# Ecological origins of perceptual grouping principles in the auditory system

**DOI:** 10.1101/539635

**Authors:** Wiktor Młynarski, Josh H. McDermott

**Affiliations:** Department of Brain and Cognitive Sciences, MIT; Center for Brains, Minds and Machines, MIT; McGovern Institute for Brain Research, MIT; Program in Speech and Hearing Biosciences and Technology, Harvard University

## Abstract

Events and objects in the world must be inferred from sensory signals to support behavior. Because sensory measurements are temporally and spatially local, the estimation of an object or event can be viewed as the grouping of these measurements into representations of their common causes. Perceptual grouping is believed to reflect internalized regularities of the natural environment, yet grouping cues have traditionally been identified using informal observation, and investigated using artificial stimuli. The relationship of grouping to natural signal statistics has thus remained unclear, and additional or alternative cues remain possible. Here we derive auditory grouping cues by measuring and summarizing statistics of natural sound features. Feature co-occurrence statistics reproduced established cues but also revealed previously unappreciated grouping principles. The results suggest that auditory grouping is adapted to natural stimulus statistics, show how these statistics can reveal novel grouping phenomena, and provide a framework for studying grouping in natural signals.

## Introduction

Sensory receptors sample the world with local measurements, integrating energy over small regions of time and space. Because the objects and events on which we must base behavior are temporally and spatially extended, their inference can be viewed as the process of grouping these measurements to form representations of their underlying causes in the world. Grouping has been viewed as a fundamental function of the nervous system since the dawn of perceptual science [1, 2, 3].

Grouping mechanisms are presumed to embody the probability that sets of sensory measurements are produced by a common cause in the world [4, 5, 6]. Yet dating back to the Gestalt psychologists, grouping has most often been studied using artificial stimuli composed of discrete elements [3, 7] – arrays of dots or line segments in vision, or frequencies in sound. One challenge in relating such research to the real world is that it is often difficult to describe natural images and sounds in terms of discrete elements. As a result, grouping phenomena have been related to natural stimulus statistics in only a handful of cases where human observers have been used to label local image features [8, 9, 10, 11, 12]. Grouping research has otherwise been limited to testing intuitively plausible grouping principles that can be instantiated in hand-designed artificial stimuli.

Grouping is critical in audition, where it is believed to help solve the “cocktail party problem” – the problem of segregating a sound source of interest from concurrent sounds [7, 13, 14, 15] (Fig. 1). As in other sensory systems, auditory grouping is believed to exploit acoustic regularities of natural stimuli, such as the tendency of frequencies to be harmonically related [16, 17, 18, 19] or to share a common onset [20, 21, 22, 23, 24, 25]. But because acoustic grouping cues have traditionally been identified using informal observation and investigated using simple synthetic stimuli, much remains unknown. First, the extent to which known principles of perceptual grouping are related natural stimulus statistics is unclear. Second, because the science of grouping has thus far been largely driven by human intuition, additional or alternative grouping principles remain a possibility.

**Figure 1:**
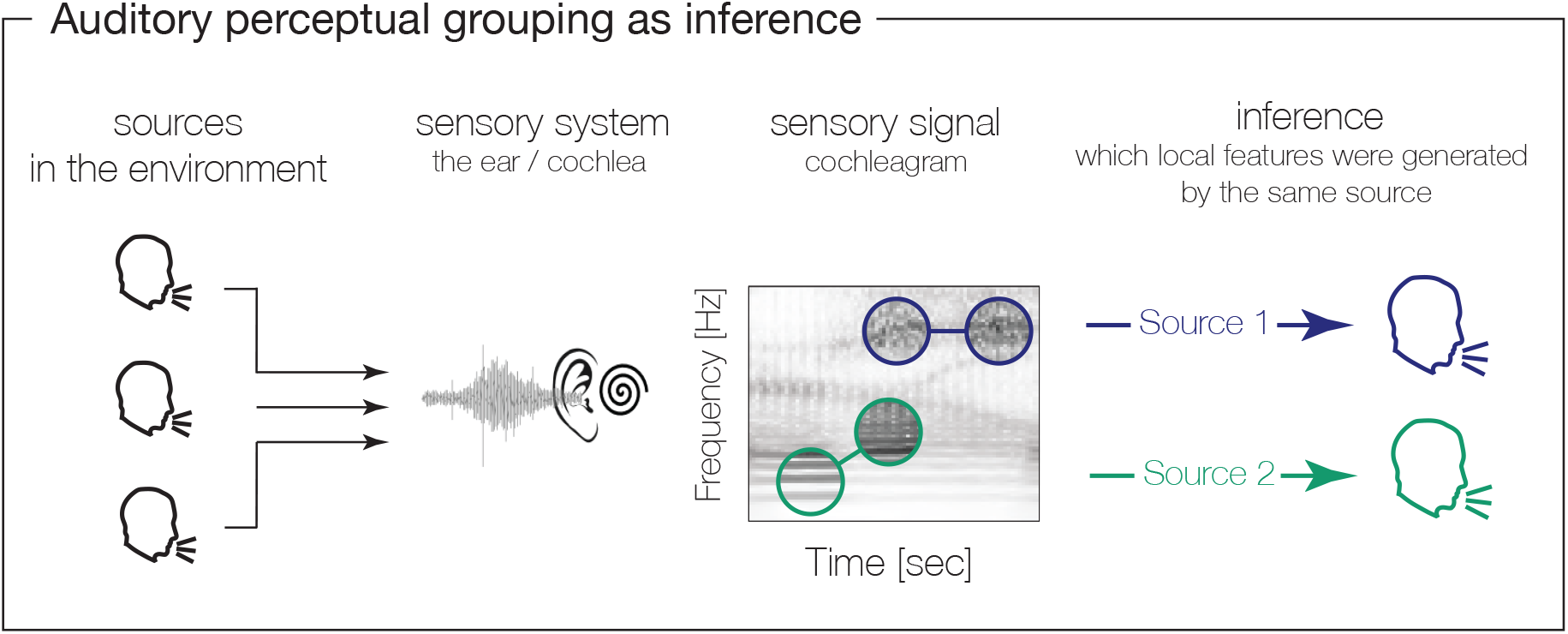
Auditory perceptual grouping. Multiple sources in the world generate signals that sum at the ear. Local sensory measurements must then be grouped to form source inferences.

We sought to link auditory grouping principles to the structure of natural sounds by measuring feature co-occurrences in natural signals and assessing their relation to perception. Unlike previous work, our approach was independent of prior hypotheses about the underlying features or the regularities that might relate to grouping. We first derived a set of primitive auditory patterns by learning a dictionary of spectrotemporal features from a corpus of natural sounds, using sparse convolutional coding [26, 27]. We then measured co-occurrence statistics for these features in natural sounds. We found that superpositions of naturally co-occurring features were more likely to be recognized as a single source than pairs of features that do not commonly co-occur, indicating that the auditory system has internalized the co-occurrence statistics over evolution or development. We next developed a method to summarize the observed co-occurrence statistics with a set of cues. The cues were instantiated as linear templates that defined stimulus properties predictive of whether features were likely to co-occur. The learned templates captured traditional grouping cues such as harmonicity and common onset, but also revealed novel grouping principles. Our results suggest that auditory grouping cues are adapted to natural stimulus statistics, and that considering these statistics can reveal previously unappreciated aspects of grouping.

## Results

In order to study grouping in natural sound signals without relying on a prior hypothesis of the features or principles that would be involved, we used convolutional sparse coding [26, 27] to first learn a set of features from which natural sounds can be composed. These features were learned from recordings of single sources of speech or musical instruments represented as ‘cochleagrams’ – time-frequency decompositions intended to approximate the representation of sound in the human cochlea. Examples of spectrotemporal features and stimuli used in all experiments can be found on the accompanying webpage: http://mcdermottlab.mit.edu/grouping_statistics/index.html.

The spectrotemporal features were optimized to reconstruct the training corpus given the constraints of non-negativity (on both feature kernels and their coefficients) and sparsity (on the coefficients). These constraints produce features that can be thought of as “parts” of the cochleagram, similar to non-negative representations of natural images [28]. The learned features capture simple and local time-frequency patterns, including single frequencies, sets of harmonic frequencies, clicks, and noise bursts (Fig. 2A), loosely analogous to the spectrotemporal features that might be detected in early stages of the auditory system [29]. Each feature can itself be viewed as an initial elementary stage of grouping sound energy likely to be due to a single source. But because natural sound signals are represented with many such features (as a set of time-varying, sparsely-activated coefficients, Fig. 2B), these features must in turn be grouped in order to estimate sound sources from the feature representation.

**Figure 2:**
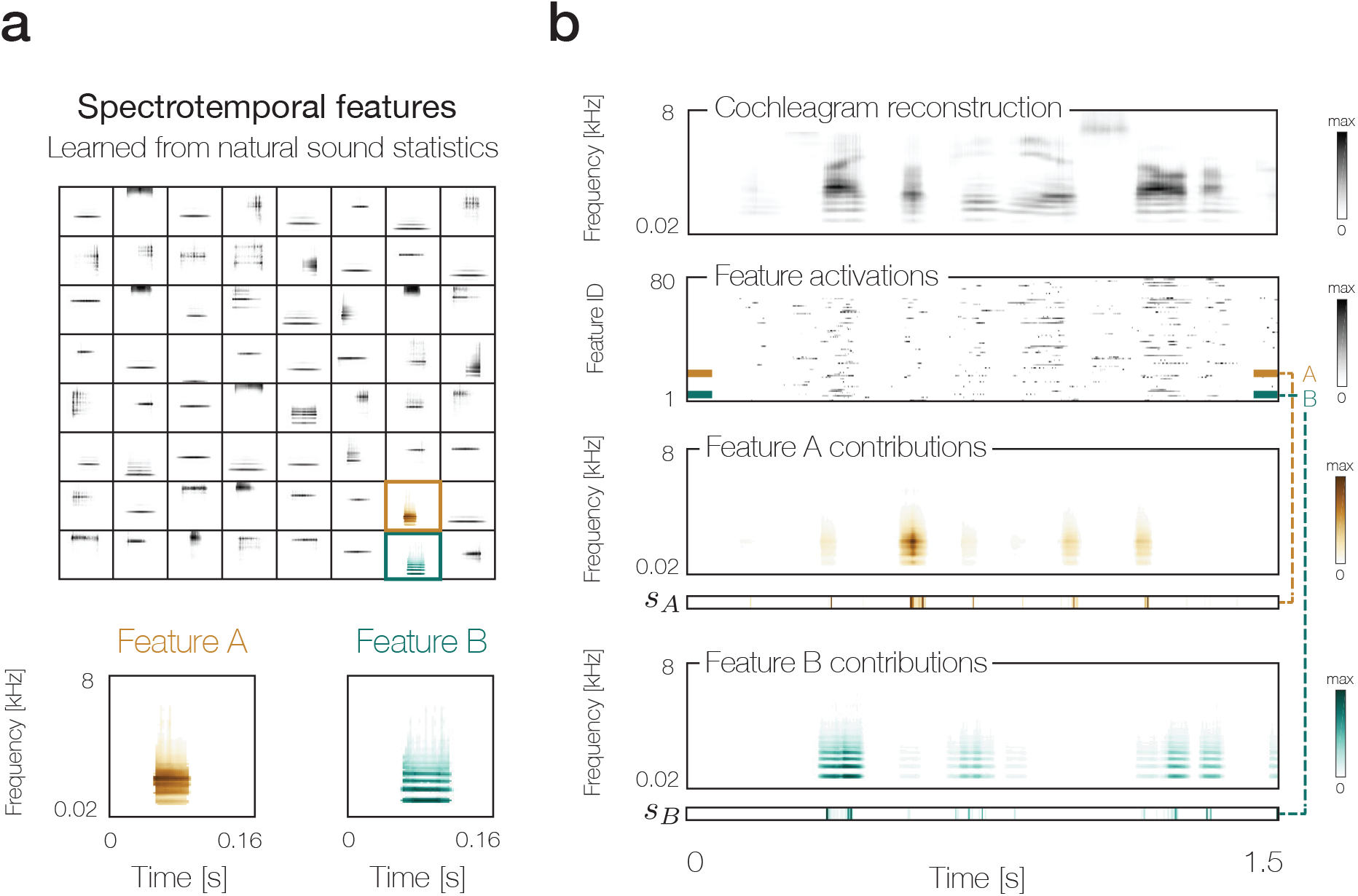
Spectrotemporal feature decompositions of natural sounds. A. Spectrotemporal features optimized to reconstruct the corpus of speech and instrument sounds. Two example features are shown at higher resolution. **B** Example reconstruction of a speech excerpt (top) from time-varying feature coefficients (second from top). The contribution of the two example features from A to the reconstruction are shown in the bottom two rows. The cochleagrams show the convolution of the feature kernel with its time-varying coefficient (shown below each cochleagram).

### Feature co-occurrence statistics in natural sounds

Once a signal is decomposed into a feature representation, the problem of grouping thus consists of determining which features are activated by the same source – an inherently ill-posed inference problem (Fig. 1). We measured co-occurrence statistics that should constrain this inference. In principle the inference of sources from feature activations could be constrained by the full joint distribution of all features. In practice this distribution is challenging to learn and to analyze [27]. Instead, we measured dependencies between pairs of features, which are tractable to measure and analyze, and which we found to contain rich structure. The key idea was to compare the co-occurrence of features within the same source to the co-occurrence of features in different sources, on the grounds that feature activations should be grouped together if they co-occur in a particular configuration substantially more often in the same source than otherwise.

To measure co-occurrences for features in the same source, we took encodings of large corpora of single sources – speech and instrument sounds – and for each feature *f* (e.g. Fig. 3A) computed the average activations of all other features at each of a set of time offsets, conditioned on the activation of the feature *f* being high (exceeding the 95th percentile of its distribution of activations) (Fig. 3B). This co-activation measure is high for features that tend to be activated at a particular time offset when the selected feature *f* is activated. To measure co-occurrence for features in distinct sources, we assumed distinct sound sources in the world to be independent. Given that assumption, the distribution of activations of one feature conditioned on the activation of another can be approximated by its marginal distribution (Fig. 3C). Thus, as a summary measure of the co-occurrence of one feature and another, we computed the ratio of the mean conditional activation of the feature to its mean marginal activation (the mean feature activation, averaged over time and across the entire training corpus). Dividing by the mean marginal activation can also be viewed as a normalization step that prevents the resulting measure from being dominated by how often a feature occurs in the training corpus. In all subsequent analyses we display the logarithm of this ratio, which we term the “association strength”. We consider a feature as co-occurring or not with the selected feature depending on whether the association strength is positive or negative.

**Figure 3:**
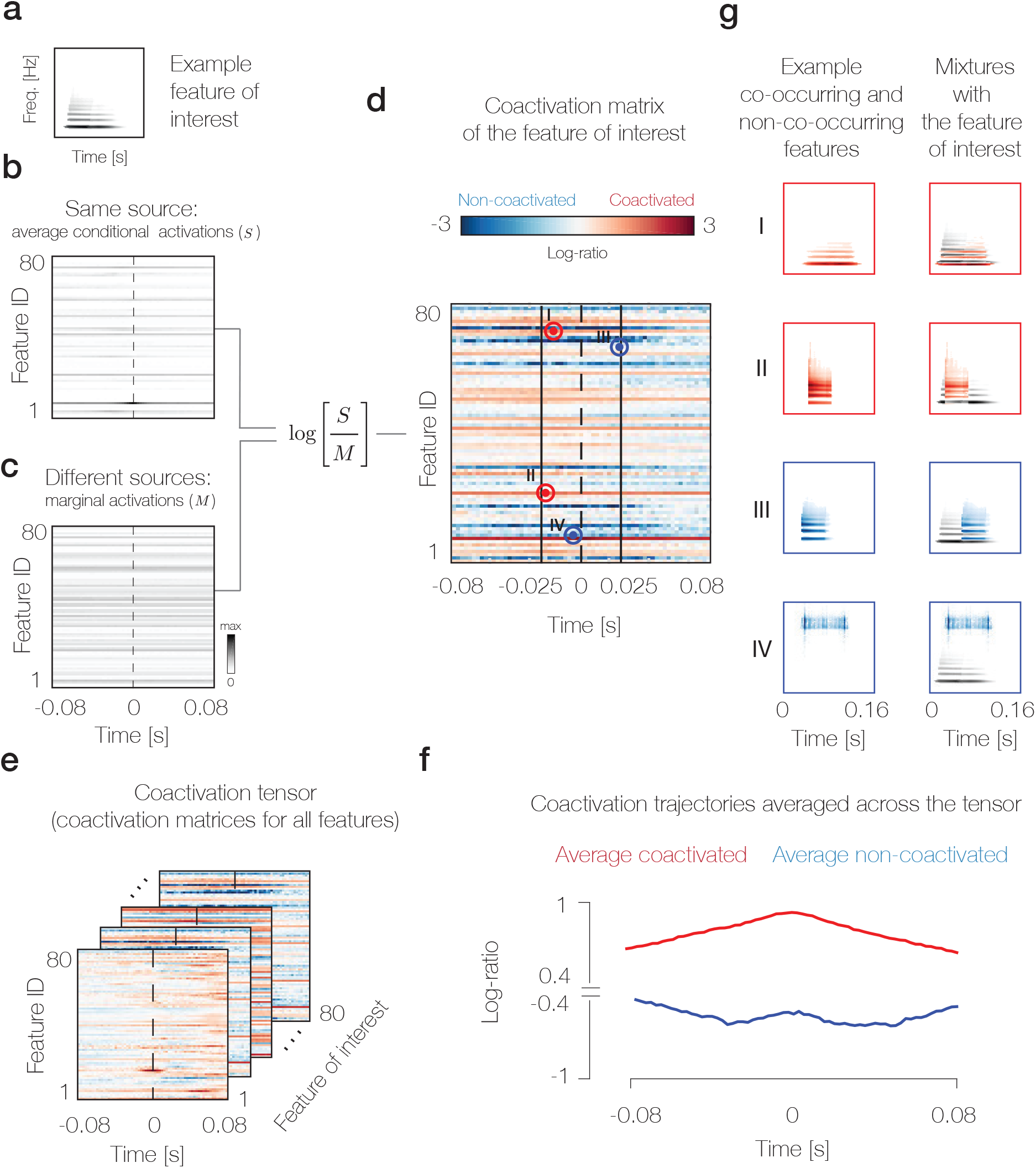
Co-occurrence statistics of spectrotemporal features. A. Example feature of interest. **B** Average conditional activations of other features, conditioned on the example feature of interest exceeding an activation threshold. **C** Average marginal activations of other features (averaged over time and across the corpus). These are by definition constant over time. **D** Co-activation matrix for the feature of interest, formed by the logarithm of the ratio of the mean conditional and marginal activations of the other features. **E** Co-activation tensor formed from the co-activation matrices of all features. **F** Positive and negative tensor entries averaged across features. The strength of association between features decreases with their time offset, as expected. **G** Examples of features with high and low association strength with the feature from A. Left column: Features colored red/blue have high/low association strength with the example feature of interest from A. Right column: mixtures of the selected features with the example feature of interest from A.

This analysis yielded a matrix for each feature (containing its association strength with each other feature at each of a range of time offsets; Fig. 3D), and thus a three dimensional tensor for the entire dictionary (Fig. 3E). These matrices are not obviously structured when inspected visually, apart from containing dependencies that on average grow weaker as the time offset between features increases (Fig. 3F). However, the tensor can be used to draw pairs of features that are strongly co-activated in the training corpus, or not, and these exhibit intuitively sensible relationships. The examples in Fig. 3G for a harmonic feature reveal that other features that strongly co-occur with it share a common fundamental frequency (f0) and fall in the same general frequency range. Conversely, features that are unlikely to co-occur with the example harmonic feature are those that are misaligned in f0 or that are far apart in frequency. These examples suggest that the co-occurrence statistics can capture reasonable relations between features, but it was not obvious to what extent the full co-occurrence tensor would have been internalized by human listeners, and to what extent it would contain comprehensible structure.

### Perceptual grouping reflects co-occurrence statistics

To test whether human listeners have internalized the measured co-occurrence statistics, we conducted a psychophysical experiment with stimuli generated by superimposing sets of features. On each trial, participants heard two such stimuli and judged which of them contained two sound sources (Fig. 4A). One feature pair was selected from the feature pairs with an association strength in the top 1% of all co-occurring pairs, and the other from the feature pairs with an association strength in the lowest 5% of the non-co-occurring pairs, i.e. the most negative (Fig. 4B; the inclusion thresholds were asymmetric because the distribution of associations strengths was asymmetric about 0). To set a ceiling level on task performance, in a second condition, one stimulus was an excerpt of a single speech or instrument sound while the other was a mixture of two such excerpts. Because natural sounds contain a superset of the dependencies measured in the co-occurrence tensor, performance on this condition should provide an upper limit on performance for the task with feature superpositions.

**Figure 4:**
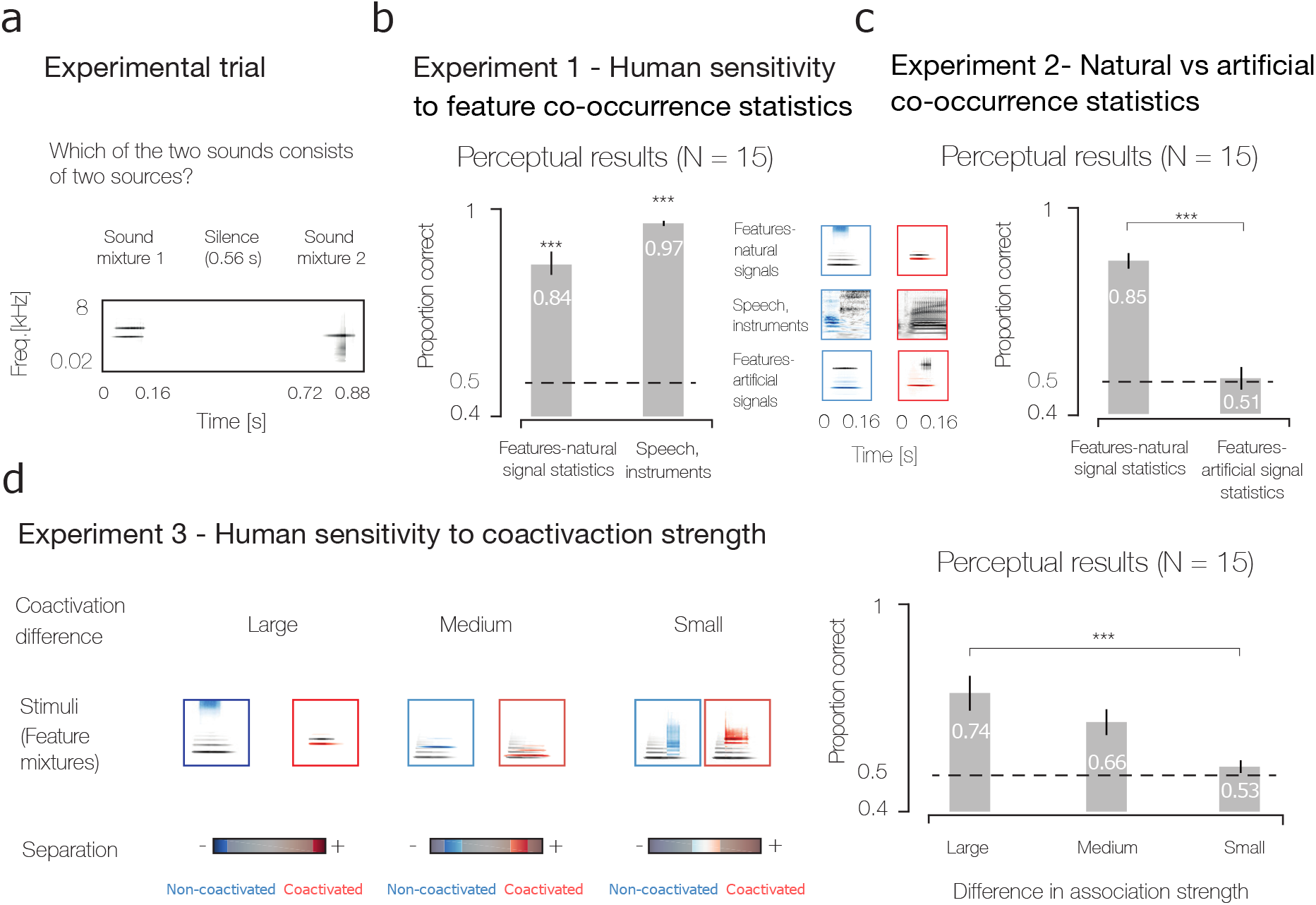
Perceptual sensitivity to natural feature co-occurrence statistics. A. Stimulus from an example trial. Listeners heard two feature pairs and judged which consisted of two sources. **B** Conditions and results of Experiment 1. Listeners discriminated i) feature pairs assembled using natural co-occurrence statistics or ii) mixtures from single excerpts of speech and/or instruments. Asterisks denote statistical significance of t-tests (vs. chance; ***: p<.001). **C** Results of Experiment 2. Listeners discriminated feature pairs assembled using i) natural co-occurrence statistics or ii) co-occurrence statistics measured from artificial sound textures. The textures were synthesized to match some of the statistics of speech (related to power and modulation spectra). Asterisks denote statistical significance of t-tests (vs chance or between conditions; ***: p<.001). **D** Conditions and results of Experiment 3. Listeners discriminated feature pairs drawn from different ranges of the co-activation continuum, producing large, medium, or small co-activation differences between the two pairs presented on a trial. Asterisks denote statistical significance of repeated measures ANOVA comparing performance in the three conditions.

Human listeners reliably identified unlikely combinations of features as sounds consisting of two sources (Fig. 4B, left bar; t-test, t(14) = 10.95, p < 0.001), only slightly below the level for speech mixtures (Fig. 4B, right bar, t(14) = 93.75, p < 0.001). This result suggests that humans have internalized aspects of the co-occurrence tensor and use the learned statistics for perceptual grouping.

To assess the extent to which the perceptual sensitivity was specific to natural co-occurrence statistics, we ran a control experiment using stimuli derived from a co-occurrence tensor measured from synthetic sound textures [30]. The textures were synthesized to match the power spectrum and modulation spectrum of speech, but were otherwise unstructured (see Methods). Co-occurrence statistics were measured using the same features learned from the natural sound corpus. The experiment thus controlled for the possibility that the features and their encoding process might themselves create dependencies that could support task performance, independent of natural signal statistics. In contrast to stimuli from the natural co-occurrence tensor, the control stimuli produced near-chance performance (Fig. 4C, right bar; not significantly different from chance, t(14) = 0.1748, p = 0.86 and significantly worse than the natural stimuli, t(14) = 10.57, p < 0.001). The results suggest that grouping judgments depend on internalized statistics that are to some extent specific to natural sounds.

To further probe the extent to which perceptual grouping judgments would reflect natural co-occurrence statistics, we generated pairs of feature pairs whose association strength difference fell into one of three ranges (Fig. 4D; each range differed from that used in Experiment 1). If listeners have internalized natural feature co-occurrences, performance should scale with the association strength difference. As shown in Fig. 4D, performance was best when the association strength difference was large, and declined as it decreased, yielding a main effect of the association strength difference (F(1.39, 19.45) = 17.46, p < 0.001). This result is further consistent with the role of natural co-occurrence statistics in perceptual grouping judgments.

### Predicting grouping cue strength from natural statistics

Grouping is typically conceptualized in terms of cues – stimulus properties that are predictive of grouping and that could thus help to solve the inference problem at the heart of grouping. We sought to relate grouping cues to co-occurrence statistics, both to evaluate the statistical validity of traditionally proposed cues and to learn cues *de novo* from statistics. We formalized a grouping cue as a function of two stimulus features whose value depends on whether the two features are likely to belong to the same source or not (Fig. 5A). We quantified the statistical strength of a cue using the co-occurrence tensor, measuring the cue for all pairs of strongly positively associated features and all pairs of strongly negatively associated features, and then quantifying the difference in the distributions of cue values for the two sets (Fig. 5B).

**Figure 5:**
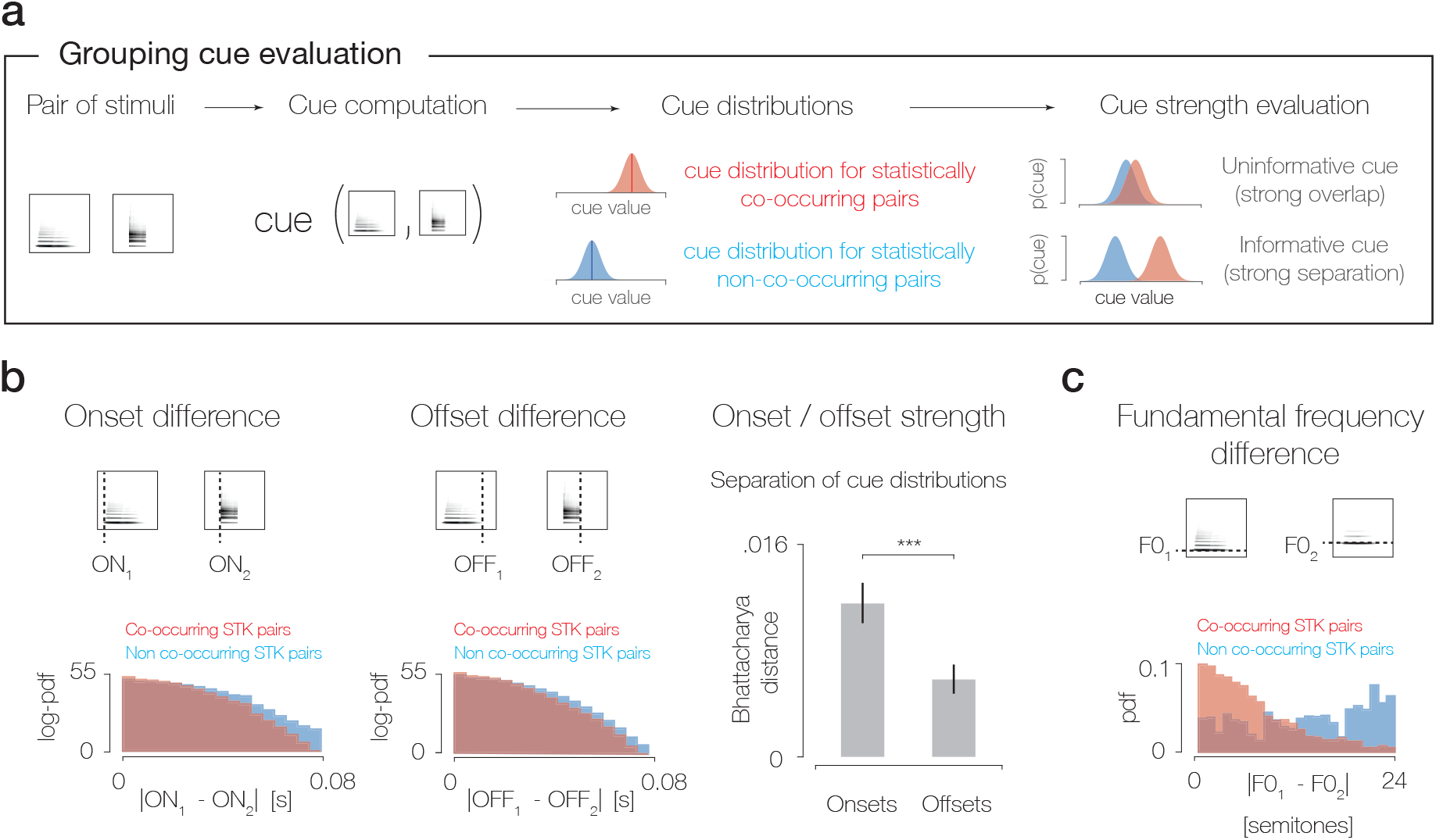
Grouping cue evaluation. A. Cues are defined as functions of feature pairs that should differ depending on whether the features are likely to be due to the same source. Cue strength is quantified as the separation of cue distributions for co-occurring and non-co-occurring feature pairs. **B** Evaluation of the cue strength of common onset and common offset. Left and middle panels illustrate cue measurement for an example feature pair (top) and the resulting cue distributions for co-occurring and non-co-occurring feature pairs (the top 75% of positive and bottom 75% of negative of feature pairs when ranked according to their association strength). Logarithmic axis serves to reveal the difference between the tails of the distributions. Right panel plots the Bhattacharya distance, a summary measure of the separation of the cue distributions for co-occurring and non-co-occurring feature pairs, predicting that common onset should be a stronger grouping cue than common offset. **C** Evaluation of the cue provided by differences in fundamental frequency (f0), which is small for co-occurring feature pairs and large for non-co-occurring pairs. This analysis was restricted to features that were above a criterion level of periodicity, and that thus had a well-defined f0.

We first considered the two most commonly cited cues from traditional accounts of auditory grouping: common onset and offset [20, 21, 22, 23, 24, 25], and common fundamental frequency [16, 17, 18, 19]. We measured onsets and offsets of each feature as the time points where their broadband envelope exceeded or dropped below a threshold value, and measured the difference in onset or offset time for all pairs of strongly positively or negatively co-activated features (corresponding to the top 75% of positive entries and bottom 75% of negative entries in the association strength tensor). Both onset and offset differences were smaller for co-activated features, but the difference was larger for onsets than offsets. This difference provides an explanation for the documented difference in the perceptual effect of common onset and offset (whereby grouping from offsets is weaker than grouping from onsets) [22]. Similarly, the f0 difference between features was smaller for co-activated features (measured in features that exceeded a criterion level of periodicity, such that the f0 was well-defined). These analyses provide what to our knowledge is the first evidence that conventionally cited grouping cues have a sound basis in natural signal statistics.

### Grouping cues derived by summarizing co-occurrence statistics

We next sought to derive grouping cues from the co-occurrence tensor in order to explore the cues that would emerge independent of human intuition. We searched for acoustic properties that would predict the association strength of feature pairs, restricting the properties to those defined by linear templates in order to facilitate their interpretability. The resulting discriminative model learned templates in the time-frequency and modulation domains whose dot-product with a spectrotemporal feature kernel was similar for frequently co-occurring features, but different for non-co-occurring features. Specifically, the model used the magnitude of the difference in the projections for two features (the ‘cue value’) to predict whether they have high association strength or not (via logistic regression; Fig. 6A). The two domains considered (time-frequency and modulation planes) are the most common representations in which to examine sound; the modulation plane is simply the two-dimensional power spectrum of the time-frequency representation of a sound [31]. Templates were learned sequentially until performance reached an asymptote, resulting in four templates, two in each of the time-frequency and modulation planes (Fig. 6B-E; additional templates only marginally improved performance, see Methods). Despite the limitations inherent to linear templates, the four templates were sufficient to differentiate co-occurring from non-co-occurring features with reasonable accuracy (81%), indicating that they captured a substantial amount of the variance in feature co-occurrence.

**Figure 6:**
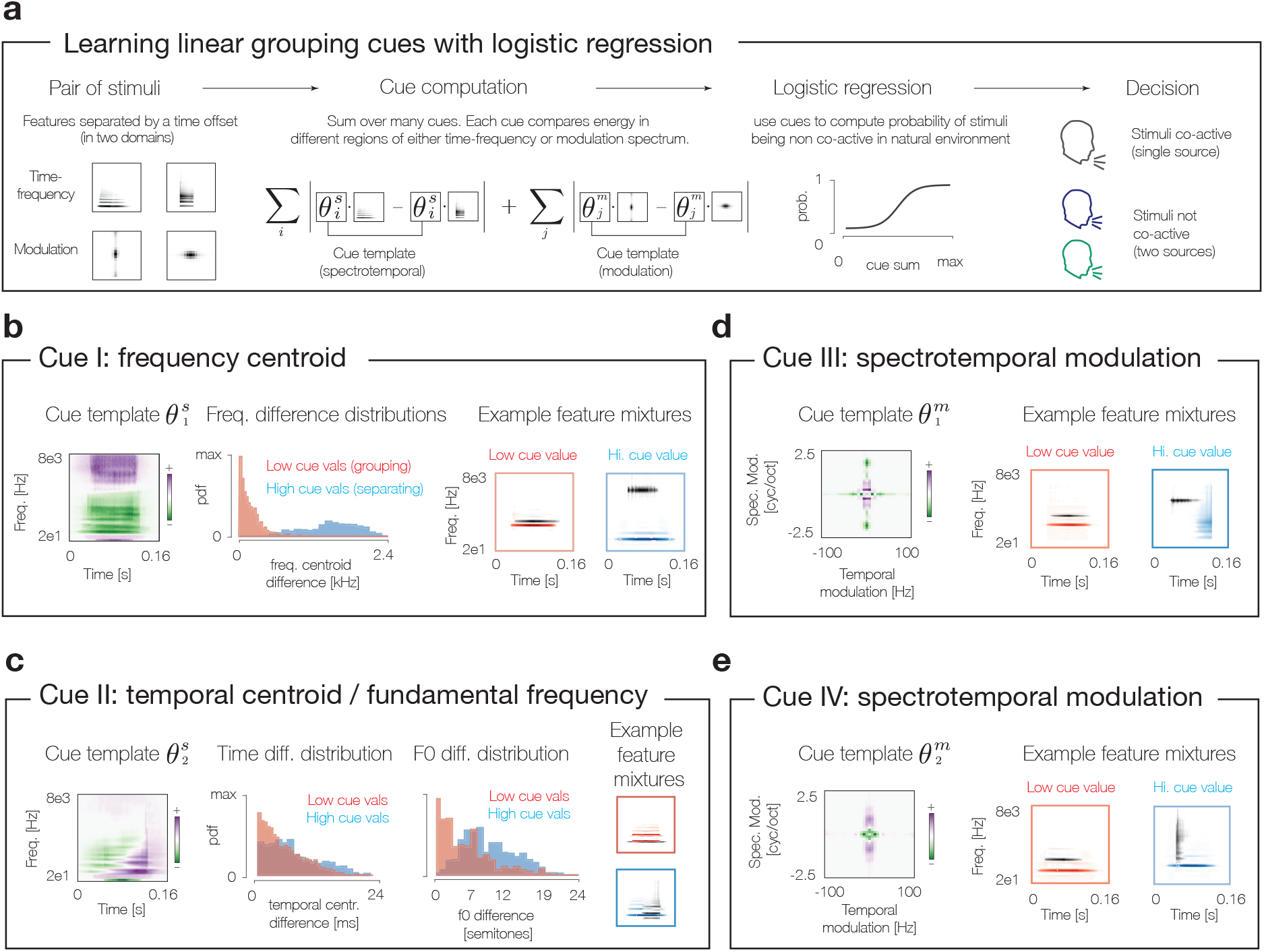
Learning grouping cues from natural signal statistics. A. Schematic of discriminative model from which cues were learned. Cues are computed for pairs of features, by projecting each feature onto a cue template and taking the absolute value of the difference. The discriminative model takes the sum of these absolute differences for a set of cue templates and predicts whether the feature pair co-occurs or not using logistic regression. The templates could be defined in either the time-frequency or modulation planes. **B-E** Characterization of the four learned cues. Left: Cue templates. Middle (B&C only): distribution of stimulus properties hypothesized to be captured by template. Right: Example feature pairs with high and low cue values. The cue in C appears to capture two conceptually distinct sound properties (temporal offset and fundamental frequency difference) with a single template.

Even though the templates were derived purely from co-occurrence statistics, without regard for prior hypotheses or human intuition, inspection of the learned templates reveals interpretable structure. The first cue template (Fig. 6B, left) can be interpreted as computing a spectral centroid, implying that features with similar frequency content are likely to co-occur. We quantified this effect by measuring the spectral centroid of each feature and comparing the centroid difference for feature pairs with high and low cue values (Fig. 6B, middle and right). Spectral differences are known to influence the grouping of sounds across time [7, 32], but this result suggests that they also should affect the grouping of concurrent sound energy (because the temporal extent of the tensor was -/+ 80 ms from the center of the reference feature, and the width of feature kernels was 162 ms, all feature pairs considered in this analysis overlapped in time to a fair extent).

The second template (Fig. 6C, right) appears to compute a temporal derivative - features that have similar projections tend to be aligned in time (Fig. 6C, middle left), recapitulating the established grouping cue of common onset/offset [20, 21, 22, 23, 24, 25]. This template also detects misalignments in fundamental frequency (Fig. 6C, middle right), another established grouping cue [16, 17, 18, 19, 33].

The modulation plane templates (Fig. 6D&E) compute differences between the power in different regions of the modulation plane, and thus capture the tendency of features with different spectral shapes (tone vs. clicks, for example) to belong to distinct sources, regardless of their temporal configuration. To our knowledge this type of cue has not been previously noted in the auditory scene analysis literature, although modulation rate has been shown to affect the grouping of sequences of sound elements [34].

### Perceptual test of learned grouping cues

The derived cues embodied in the templates varied in their statistical cue strength, but all were individually predictive of whether feature pairs were associated or not (Fig. 7A, using the analysis of Fig. 5). To test whether the derived cues affect perceptual grouping, we used each individual template to construct experimental stimuli, and measured whether listeners’ ability to use the cue in a grouping judgment varied in accordance with its statistical strength in the training corpus of natural sounds. For each cue, we searched for pairs of features with high values of that cue but low values of the other three cues, such that the cue of interest would provide the only indication that the two features were not part of the same source (Fig. 7B, left). We then presented the pair successively with another pair in which all four cues had low values, and asked listeners to judge which of the two pairs consisted of two sources. Listeners were significantly above chance for each cue (Fig. 7B, right; t(14) ≥4.17, p<.001 in all cases), suggesting that all cues contribute to perceptual grouping judgments. Moreover, performance varied with the statistical cue strength, providing additional evidence that perceptual grouping is based on internalized co-occurrence statistics.

**Figure 7:**
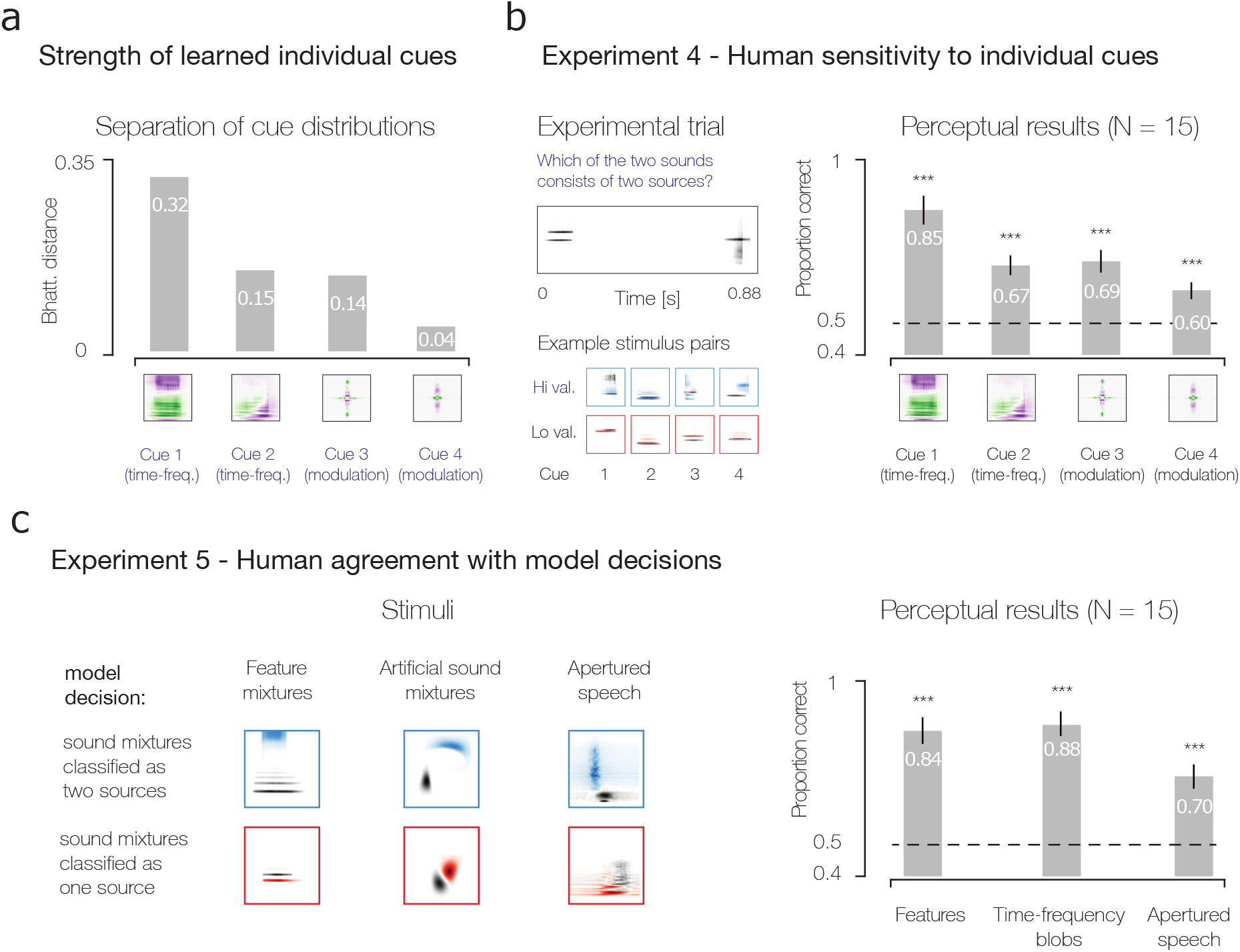
Perceptual sensitivity to learned grouping cues. A. Cue strength of the learned cues, measured as the Bhattarcharya distance between the cue distributions for co-occurring and non-co-occurring feature pairs. **B** Description and results of Experiment 4, which measured perceptual sensitivity to each of the four learned cues. The task was the same as in Experiments 1-3: listeners heard two feature pairs and judged which one consisted of two sources. One feature pair on a trial had a low cue value (implying high association strength) and one had a high cue value (implying low association strength). **C** Description and results of Experiment 5, which measured human agreement with model decisions about the segregation of mixtures of three types of stimuli (the spectrotemporal kernels learned from speech and instruments, artificial time-frequency “blobs”, and speech excerpts windowed in the time-frequency plane). On each trial listeners heard two mixtures and judged which consisted of two sources.

As a further test of the predictive value of the learned cues, we used them to predict the perceptual grouping of three types of stimuli: pairs of the learned spectrotemporal kernels, mixtures of artificial sounds synthesized from “blobs” in the time-frequency plane, and mixtures of speech segments win-dowed by time-frequency apertures. Apertures were used for the speech conditions because mixtures of extended speech excerpts almost never perceptually group to resemble a single source. We searched for stimuli that the cue model confidently judged to be single sources as well as stimuli that the model confidently judged to be mixtures, and on each trial presented listeners with one stimulus from each group, asking them to identify the single source. In all three cases listeners’ judgments agreed with those of the model (being well above chance for each condition; t(14)*≥*5.82, p<.001 in all cases). These results provide further evidence for the perceptual reality of the derived cues, and show that they have fairly general predictive power.

### Grouping of feature sequences

Experiments 1-5 demonstrate the perceptual relevance of empirical co-occurrence statistics, and of the cues that we derived from them, but utilized pairs of features or sound excerpts in close temporal proximity. To test whether the measured co-occurrence statistics would be predictive of the perceptual grouping of more extended sound sequences, we used the co-occurrence tensor to generate sequences of features spaced more widely in time, and measured whether the co-occurrence statistics could predict the perceptual “streaming” of these sequences. Each sequence was seeded with an initial feature. Subsequent features were chosen from a probability distribution derived from their association strength with the previous feature, with features with higher association strength having higher probability (Fig. 8A). For each of a set of reference sequences, we generated two types of mixtures: one with a second sequence whose features had high association strength with the features of the reference sequence,and one whose features did not (Fig. 8B; see Methods). Listeners were presented with a mixture and judged whether it was generated by one or two sources.

**Figure 8:**
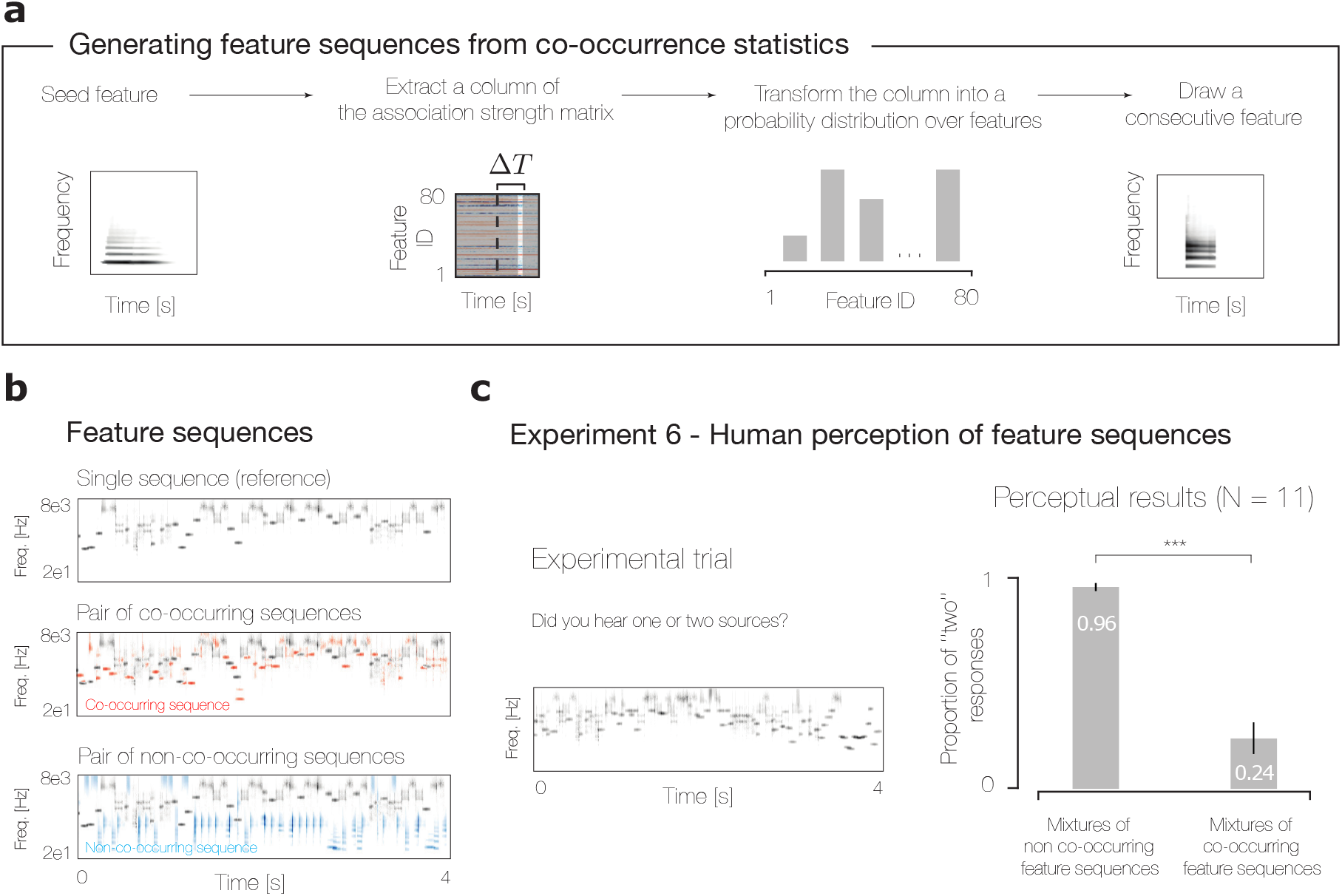
Streaming of spectrotemporal feature sequences. A. Sequence generation from co-occurrence statistics. First, a seed feature is chosen. Second, the column of its association strength matrix is extracted corresponding to the desired time offset for the next feature (here fixed at 75 ms). Third, the column is transformed to a probability distribution via the softmax function. Fourth, the next feature is drawn from this distribution. These steps are iterated until a sequence of the desired length is obtained. **B** Example reference sequence (top), mixed with a co-occurring sequence (middle), and non-co-occurring sequence (bottom). **C** Description and results of Experiment 6. On each trial listeners heard a mixture of two feature sequences and judged whether it was produced by one or two sources. Asterisks denote statistical significance of a paired t-test between conditions.

As shown in Fig. 8C, listeners reliably judged the mixture with the non-co-occurring sequence as two sources, but showed the opposite tendency for the mixtures with the co-occurring sequence (t(10) = 9.56, p <.001, t-test). Subjectively, the sequences in a non-co-occurring mixture typically differed in their acoustic qualities, and attention could often be directed to one or the other. There was thus some similarity to classical examples of streaming with alternating tones and other simple sound elements [32, 35] even though the sound sequences here were more stochastic and varied. The results indicate that pairwise co-occurrence statistics capture some of the principles that cause extended sound sequences to perceptually stream.

## Discussion

We introduced a framework for measuring natural signal statistics that could underlie perceptual grouping, and explored their relationship to perception in the domain of audition. We first learned local acoustic features from natural audio signals (Fig. 2) and computed their strength of co-occurrence (Fig. 3). Our results revealed that acoustic features exhibit rich pairwise dependencies, but also that these cooccurrences could be summarized to a large extent with a modest number of “cues”. We formalized the notion of a cue as a stimulus property that predicts the co-occurrence of pairs of features (Fig. 5), and derived cues from the large set of measured pairwise co-occurrence statistics (Fig. 6). The cues that emerged include some previously known to influence grouping, such as common onset and fundamental frequency, but also some factors that have not previously been widely acknowledged (such as separation in acoustic and modulation frequency for concurrent features). We found evidence that the auditory system has internalized these statistics and uses them to group features into coherent objects. This was true both for isolated pairs of features (Experiments 1-3) and for more extended feature “streams” (Experiment 6), and for each of the individual cues revealed by the co-occurrence statistics (Experiment 4). These results provide what to our knowledge is the first quantitative link between auditory perceptual grouping and natural sound statistics, show how these statistics may be harnessed to study auditory scene analysis, and offer a general framework for relating natural signal properties to perceptual grouping.

### Related work

The derivation of ideal observer models has a long and productive history in perception research [36], and such models have been used to learn cues for a range of natural tasks [37]. However, previous such attempts to relate perceptual grouping to natural scene statistics have largely been limited to contour grouping in images [8, 9, 10, 11, 12]. These influential earlier efforts inspired our present work, but were reliant on hand-picked features labeled by human observers (object edges), and their analysis was limited to dimensions thought to be important a priori (position and orientation). Our results demonstrate how one can derive grouping cues from features learned entirely from natural signals without prior hypotheses about the features or underlying grouping principles. Learning signal features and grouping cues from the structure of natural sounds paid dividends by revealing statistical effects that were not obvious beforehand and that were found to underlie novel perceptual effects. Our methodology also gives additional support to commonly discussed cues, by showing that they emerge from the large set of possible cues that might in principle have been derived from natural sound statistics.

Our results complement a long research tradition that has documented behavioral and neural effects of a handful of acoustic grouping cues, relying on intuitively plausible cues and synthetic stimuli [7, 16, 17, 18, 19, 20, 21, 22, 23, 38, 39, 40]. Our results provide statistical justification for the two most commonly studied cues from this literature (onset and harmonicity), but also identify other statistical effects, and show their perceptual relevance. Frequency separation is known to strongly affect the grouping of stimuli over time [32], but is less acknowledged to influence the grouping of concurrent features. Our results show that it is the strongest effect evident in local co-occurrence statistics of natural audio, at least for the corpora that we analyzed, and that it has a correspondingly strong perceptual effect. Modulation separation has also not been appreciated as influence on the grouping of concurrent features [34], but emerged from the analysis of co-occurrence statistics and also proved to have a large perceptual effect. The analysis of natural signal statistics is thus both “post-dictive”, suggesting normative explanations for known effects, but can also be predictive, pointing us to phenomena we should test experimentally.

Our quantitative approach to grouping has the added benefit of taking us beyond verbal descriptions of phenomena to enable grouping predictions for arbitrary stimuli. We leveraged this ability to make such predictions for three different types of stimuli (Experiment 5). The verbal characterizations of cues from classical approaches cannot be tested in this way.

Our approach also complements engineering efforts to solve auditory grouping. Early attempts in this domain were inspired by psychoacoustic observations, and thus implemented hand-engineered grouping constraints based on common onset and periodicity [41, 42, 43]. More recent attempts to build computational models of sound segregation similarly focus on the intuitively plausible cue of temporal coincidence [24, 25]. Current state-of-the-art engineering methods instead rely on learning how to group acoustic energy from labeled sound mixtures [44, 45], but are at present difficult to probe for insight into the underlying acoustic dependencies. Our methodology falls between these two traditions, utilizing the rich set of constraints imposed by natural signals but providing interpretable insight into factors that might underlie grouping. Indeed, our choices to restrict the analysis to pairwise dependencies, and to learn linear cues that summarize the measured dependencies, were made to facilitate inspection of the results. It could be fruitful to apply our analysis methods to contemporary source separation methods, as well as to use the dependencies that we measured to separate sources from mixtures.

### Open issues and future directions

Our approach leveraged available recordings of single sound sources. Single source recordings provide a weak form of supervision, in that the resulting feature activations can be assumed to belong together without requiring the use of human labels that were critical to previous work in this vein [8, 9, 10, 11, 12]. However, because large numbers of single source recordings are presently available only for speech and musical instruments, our analysis was limited to these sound genres, and as such may not be representative of the full set of dependencies in natural sounds. The use of available audio recordings had the additional consequence that our analysis was restricted to monaural audio. Natural auditory input likely contains important binaural dependencies that contribute to grouping [40, 46, 47, 48, 49], that our approach could in principle capture if applied to audio recorded from two ears [50]. Another limitation of our approach lies in the use of sparse feature decodings, which efficiently describe speech and music sounds, but are a poor description of more noise-like sounds such as textures. Textures are an important part of auditory scene analysis [51], and studying the statistical basis of their grouping will likely require an alternative encoding scheme, potentially based on summary statistics [52] rather than localized time-frequency features.

Our results suggest that human listeners have internalized the co-occurrence statistics that we measured, in that listeners reliably discriminate between feature pairs with high and low association strength (Fig. 4). However, the results leave open whether knowledge of the dependencies is built into the auditory system over evolution, whether it is learned during development, and/or whether it continues to be updated during adulthood. Some types of sound source structure can be learned relatively quickly and can aid source separation [54], but it remains unclear whether such rapid learning effects apply to the sorts of local feature dependencies studied here. This could in principle be addressed by exposing listeners to sounds with altered statistical dependencies and then measuring whether perceptual grouping is altered.

A full account of auditory scene analysis will undoubtedly require more complete statistical models of natural sound sources, incorporating more than the pairwise dependencies between local features studied here [27]. In addition to multi-element dependencies, a full model will likely require additional hierarchical structure, in which groups of local features are in turn grouped into larger-scale configurations. Such hierarchical organization could be one way to model the grouping effects of repetition [55, 56], which is one powerful grouping phenomenon not accounted for by our analysis.

The instantiation of perceptual grouping in the brain remains a key open issue in systems neuroscience, particularly in audition [24, 57, 58, 59]. The features that we measured could plausibly be detected by neurons in the auditory system [29], and the co-occurrence statistics that we analyzed could in principle be encoded by connections between neurons representing local features, analogous to the association fields for contour grouping which are thought to be instantiated in lateral connections between visual neurons [60]. Alternatively, co-occurrence statistics could be encoded by higher-level sensory neurons implementing logical AND/OR-like computations [61, 62, 63]. The latter possibility could be tested by comparing the components of such multi-dimensional receptive fields to the cue templates that we derived.

Although our methodology starts from an encoding scheme based on local features, in part because these are most readily mapped onto early stages of sensory systems [64, 65, 66], problems of scene analysis can also be approached with generative models more rooted in how sounds are produced. For instance, speech and instrument sounds are fruitfully characterized as the product of a source and a filter that each vary over time in particular ways [67, 68], as are sounds in reverberant environments [69], and humans appear to have implicit knowledge of this generative structure [70]. Reconciling these generative models for sound with those rooted in neurally plausible local feature decompositions is a critical topic for future research.

## Materials and Methods

### Natural sound corpus

We created a corpus of sounds generated by individual physical sources by merging corpora of recordings of individual talkers and musical instruments in equal proportion. Speech sounds were taken from the TIMIT database [71], and included voices of male and female speakers speaking sentences in English. Solo instrument sounds were taken from the RWC Music Database72. The database consists of individual notes played by a diverse set of instruments including pianos, guitars, brass, woodwinds and drums. We uniformly sampled random excerpts of sound from all recordings in the database. The final dataset consisted of 7000 excerpts (3500 excerpts of speech and 3500 excerpts of instruments), each 3 seconds long, resulting in approximately 5 hrs 48 minutes of sound. The sampling rate was set to 16 kHz.

### Cochleagrams

All analyses used a cochleagram representation of sounds intended to approximately simulate the output of the auditory nerve. Cochleagrams were generated as in previous publications [52, 30]. Raw sound waveforms were passed through a bank of 81 bandpass filters. Filters were regularly spaced on an equivalent rectangular bandwidth (ERB) scale with bandwidths matched to those expected in the healthy human ear [72]. Center frequencies spanned 31 Hz - 7656 Hz. Transfer functions were shaped as the positive portion of a cosine function. The cochleagram was generated from Hilbert envelopes of the output of each filter, transformed with a power function (with the exponent 0.3, roughly approximating properties of the basilar membrane compression [73]). The result was downsampled to 400 Hz.

### Learning the feature dictionary

To learn an acoustic feature basis for cochleagrams we used a convolutional sparse-coding model described in [27] with an additional non-negativity constraint imposed on the basis functions, to aid interpretability in terms of sound energy (which is non-negative) and produce localized features [28]. The model represents a cochleagram excerpt as a sum of spectrotemporal kernels (STKs) *ϕ* (162 ms in duration) convolved with their activation time courses *s*:

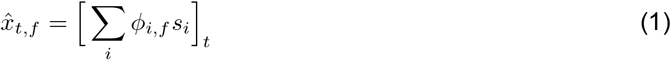

The model finds feature activations for individual cochleagram excerpts by minimizing the following cost function:

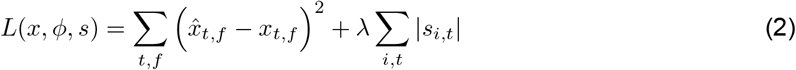

where λ is a parameter controlling the degree of sparsity. The sparsity term in equation 2 implicitly assumes that feature activations follow an exponential distribution.

A feature dictionary was learned from the speech/instrument corpus described above with the following standard iterative two-step learning procedure. All spectrotemporal kernels were first initialized with Gaussian noise. During each learning epoch a random cochleagram excerpt (320 ms in length, i.e. 129 samples) was drawn from the dataset. In the first step, optimal coefficients were inferred for the cochleagram excerpt by minimizing equation 2 with respect to sparse coefficients s. In the second step, the inferred coefficients were used to perform a gradient step on the basis functions ϕ. The two steps were iterated (each time with a different randomly drawn cochleagram excerpt) for 100, 000 epochs. The value of the sparsity controlling parameter *λ* was set to 0.2.

Because the inference of all coefficients s is computationally demanding, we relied on an approximate inference scheme75. Instead of inferring the values of all coefficients for each excerpt, we selected a subset of them to be minimized. This subset consisted of the 1024 coefficients *s_i,t_* where the associated kernels *ϕ_i_* generated the strongest projections on the cochleagram (i.e. best matched the structure of the signal). During the inference process, only the values of these coefficients were optimized, while the others were set to 0. The gradients of the basis functions were then computed using the coefficients from this approximate inference step.

We set the number of learned kernels to 80. We found empirically that if this number was larger, some of the kernels would not converge during training. Because different random initializations yield slightly different sets of feature kernels, we trained 10 different sets of kernels, and then combined them for subsequent analyses as described below. The analyses were thus based on a total of 800 learned kernels.

### Co-occurrence statistics

Association strength matrices were computed by first averaging a feature’s coefficients conditioned on another feature exceeding an activation threshold. Using the learned features *ϕ*, we first inferred optimal coefficients *s_i,t_* for each of the 7000 3-second-long sound excerpts in the sound corpus. In that way we obtained 7000, 3-second long (1200 samples) coefficient maps. Rows of coefficient maps corresponds to individual kernels *ϕ_i_* and columns to time points *t*. From each coefficient map generated in that way we then sampled 50 random, 160 ms (65 samples) long excerpts. This resulted in a dataset of 350 000 excerpts of coefficient maps.

For each kernel of interest *ϕ_i_* we selected the coefficient map excerpts where the activation coefficient *s_i_* at the excerpt’s center (i.e. t=33) exceeded an activation threshold *τ_i_*. The activation threshold *τ_i_* was set to be equal to the 95th percentile of the distribution of coefficients *s_i,t_*, estimated using the entire dataset. The coefficient map excerpts selected in this way were averaged to obtain the conditional activation matrix *S*. We note that one justification for using the mean conditional activation as a measure of dependence is that the features were learned assuming an exponential prior on the coefficients, whose scale parameter is fully captured by the empirical average.

Marginal kernel activations were computed by averaging the corpus encodings across time and samples, resulting in a vector *v_i_* of average activations of each kernel *ϕ_i_*. This vector was then concatenated 65 times to create a marginal activation matrix *M* (since the marginal activation by definition does not depend on time).

Association strength matrices for each kernel *ϕ_i_* were then computed by taking the logarithm of the element-wise ratio of the corresponding conditional activation map *S* and the marginal activation map *M*. This procedure was followed for each of the 10 feature dictionaries, yielding 10 different tensors.

One interpretation of this ratio is that it compares the expected co-activation of a feature with another when they are generated by the same source vs. when they are generated by different sources. This interpretation assumes that different sources are independent, such that the distribution of a feature conditioned on another being active is just that feature’s marginal distribution. Another interpretation is that the ratio serves to normalize the conditional activation of a feature by its mean activation, so that the quantity can be compared across features that have different average activations.

### Computing cues

#### Onset/offset detection

The onset of each STK was computed from the mean across frequency channels of the subband temporal envelopes composing its cochleagram. Onset time was defined as the first time point (measured from the beginning of the kernel) at which the envelope exceeded 5% of its maximal value. Analogously, offset time was defined as the time point where the envelope dropped below the 5% threshold of the maximal value for the last time.

#### F0 extraction with YIN

Periodicity and fundamental frequency of each kernel were computed using the YIN pitch tracking algorithm [74] applied to a waveform representation of the kernel (see below for details of cochleagram-to-waveform inversion method). We analyzed F0 differences only among kernels with an aperiodicity index below 0.2.

### Stimulus generation - cochleagram inversion

Stimulus waveforms were generated from cochleagrams via an iterative inversion procedure. The waveform was initialized with white noise. Each iteration consisted of the following steps:

1. Generate subband decomposition of waveform using cochlear filterbank.
2. Divide out Hilbert envelopes of each waveform subbands and multiply by the corresponding cochleagram envelope.
3. Refilter the modified subbands and sum to yield a new waveform

These steps were repeated 20 times. Iteration was necessary because step 3 altered the subband envelopes, partially undoing the effect of step 2. Over time the resulting waveform converged to a state in which the subband envelopes were close to the desired values.

### Learning grouping cues through discriminative model training

The purpose of the discriminative model was to learn acoustic properties that were predictive of the co-occurrence of STK pairs in the training corpus. We quantified acoustic properties with linear templates in the time-frequency and modulation planes (the two most common domains in which to analyze sound). The discriminative model learned templates in the time-frequency and modulation domains (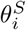and 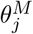 respectively) whose dot-product with an STK was similar for frequently co-occurring STKs, but different for non-co-occurring STKs. A grouping cue was thus operationalized as the absolute value of the difference in template projections between two sounds in one of the two domains:

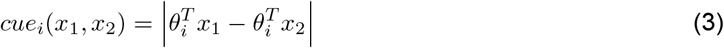

where T denotes the transposition operator. Although the model was trained using STKs, it could be applied to an arbitrary pair of sounds, which we denote *x*_1_, *x*_2_.

For each pair of kernels, represented in the spectrotemporal and modulation domains (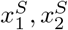 and 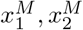 respectively), the following sum across all cues was computed:

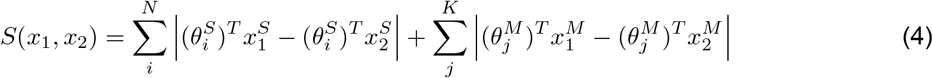

where each term in each of the sums is a value of a cue corresponding to a particular template. The probability of the two sounds being non-co-occurring in the training set was then computed by applying a logistic nonlinearity to *S*(*x*_1_, *x*_2_):

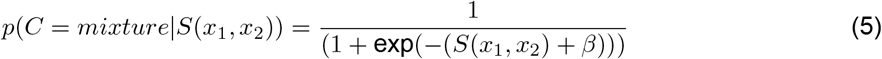

Cues were learned in a greedy fashion. First, the total desired number of cues was chosen (here, *N* + *K* = 4, chosen because this number was found empirically to produce good discrimination performance, but was not so large as to preclude inspection of individual cues). Adding additional cues only marginally improved discrimination performance (4 cues yielded 81% correct, 12 cues gave 82%, and 16 cues gave 83.5%). The sub-ceiling asymptotic performance presumably reflects limitations of linear cues, which we adopted to facilitate inspection rather than maximize discriminative performance. Nonlinear operations are likely needed to fully capture some quantities that are important for grouping, and to maximize discrimination. Nonlinear cues could in principle be explored using a similar framework, but would likely be more challenging to interpret.

During each iteration a new cue template was learned in the time-frequency domain, and another one in the modulation domain. These cue templates were learned by maximizing the log-likelihood of the data via gradient descent. In the next step, the cue template (either time-frequency-or modulation-based) that increased the data log-likelihood by the largest amount was retained and incorporated into the cue basis. The other cue was discarded. These steps were iterated until the total desired number of cues was learned. Cue templates within each domain were constrained to be mutually orthogonal.

To create the training dataset, we combined co-occurring and non-co-occurring STK pairs corresponding to positive and negative entries within the co-occurrence tensor respectively. From individual co-occurrence matrices corresponding to each STK, we selected STK pairs corresponding to the highest positive 512 entries and the lowest negative 512 entries. Since each matrix consists of 5200 entries, this approximately corresponded to the upper and lower 10% of entries within each co-occurrence matrix. To facilitate learning, templates were learned in a lower dimensional subspace. The dimensionality of the time-frequency feature representations was reduced with principle components analysis to 32 dimensions. These 32 dimensions accounted for 72% of the variance across features. The dimensionality of features in the modulation domain was reduced to 16 dimensions, accounting for 99% of variance. We experimented with different numbers of PCs and these settings produced the best convergence out of those that we tried. Once learned, the templates were projected back to the stimulus space for display purposes.

### Perceptual experiments

All experiments were approved by the Committee on the use of Humans as Experimental Subjects at the Massachusetts Institute of Technology, and were conducted with the informed consent of the participants.

#### General setting

With the exception of Experiment 6 (streaming of STK sequences), each experiment followed the same 2-AFC design. During each trial participants heard two 0.16 second-long sounds separated by a 0.5 second-long silence period. Participants were asked to judge “Which of the two sounds consisted of two different sources”, and indicated their choice on the keyboard. Participants were allowed to listen to the two stimuli as many times as they wished on each trial. All stimuli were presented at 70 dB SPL over Sennheiser HD 280 Pro headphones, played out via a Mac Mini computer. A 20 ms Hanning window was applied to the beginning and the end of each sound to prevent onset/offset transients.

#### Experiment 1 -Sensitivity to STK co-occurrence statistics

For each STK in each of the learned feature dictionaries we created co-occurring mixtures and non-co-occurring mixtures by pairing it with other STKs. Because the distribution of the association strength was both asymmetric about zero and variable in shape and extent across STKs, we used two criteria to select the STK pairings. First, the pair had to have an association strength in the top 1% of all positive association strength values (for the co-occurring mixtures) or in the bottom 5% of all negative association strength values (for the non-co-occurring mixtures). Second, each STK could contribute at most 9 mixtures of each type. These mixtures were chosen to be those that had maximal (for co-occurring mixtures) or minimal (for non-co-occurring mixtures) association strength values subject to the first constraint. Additionally, the mixtures were constrained to lie in the central temporal region of the co-activation matrix of that STK (specifically, the entries within 25 ms of the center; see Experiment 6 for stimuli with more widely spaced features). Each value corresponded to a particular STK and time offset, and the two STKs were superimposed with the designated time offset. In that way we could obtain at most 9 ×80 ×10 = 7200 co-occurring STK mixtures and 7200 non-co-occurring mixtures. The combination of the selection constraints with the empirical distributions of association strengths resulted in 7156 co-occurring STK mixtures and 2775 non-co-occurring mixtures.

On each trial one mixture (superposition) of co-occurring STKs and one of non-co-occurring STKs were selected at random. A response was considered correct if a participant selected the interval containing the non-co-occurring STKs. Participants completed 100 trials with STK mixtures derived from natural sounds statistics.

In a separate block, we tested participants’ ability to discriminate individual natural sound sources from mixtures thereof. Using the sounds used to train the STKs, we generated 100 random excerpts of individual sources (speakers and instruments) and 100 mixtures of two random excerpts (speakers and/or instruments). Each excerpt had a duration equal to that of an individual STK. Participants completed 100 trials with these natural stimuli. Performance on the odd-numbered trials was used to select participants (to eliminate participants who might have misunderstood the task, or who might not have been motivated, as described below in the Participants section), and was then discarded. Only the even-numbered trials were used for the analyses in the paper, to avoid errors of non-independence.

#### Experiment 2 - Sensitivity to coactivation strength in artificial sounds

The experiment was identical to Experiment 1 except that a condition with stimuli derived from cooccurrence statistics of a corpus of artificial sounds was substituted for the speech/instrument condition. Artificial sound textures were synthesized to match a set of statistics measured in speech. Specifically, we used the synthesis algorithm of McDermott and Simoncelli [30], imposing the marginal statistics (mean and variance) of cochlear filter envelopes and the power in each of a set of modulation filters applied to the cochlear envelopes. These statistics were chosen to create stimuli with naturalistic spectra and modulation content, so that they would be well described by the feature set learned from natural sounds, but to otherwise lack the statistical dependencies present in natural sounds. Statistics were measured and imposed using an auditory model identical to that described in the original publication [30] except that the cochlear filterbank parameters were changed to those used to generate the cochleagrams from which co-occurrence statistics were measured. We generated 600 excerpts of such textures, each 6 seconds long. Each excerpt had statistics matched to a unique, random combination of 20 sentences from the TIMIT database. Each sound was split into two 3 second excerpts. These excerpts were then encoded using the feature dictionaries learned from speech and instruments, and experimental stimuli were derived using the same procedure as for condition 1 of Experiment 1. The other experimental condition was identical to condition 1 of Experiment 1.

#### Experiment 3 - Sensitivity to coactivation strength

The experiment was identical to Experiment 1 except that there were three conditions, differentiated by the magnitude of the difference in association strength between co-occurring and non-co-occurring STK pairs. Co-occurring STK pairs were drawn from the following association strength intervals: [2, 10] (condition 1), [1, 1.2] (condition 2), [0, 0.2] (condition 3). Non-co-occurring STK pairs were drawn from the following intervals [−10, −2] (condition 1), [−1.2, −1] (condition 2), [-0.2, 0] (condition 3). These intervals were selected to approximately uniformly span the range of values of the co-activation tensor entries. In a manner analogous to Experiment 1, for each of the three conditions we generated up to 9 co-occurring stimulus mixtures and 9 non-co-occurring mixtures per STK, randomly sampled from the interval.

During the experiment participants completed 70 trials from each class (210 trials in total) in a random order.

#### Experiment 4 - Sensitivity to individual learned cues

Stimuli were selected from an initial set of 50,000 STK mixtures consisting of pairs of STKs randomly drawn from 10 learned dictionaries, at random time offsets within the [0, 50] ms range. We computed cue values for each STK mixture (the absolute value of the difference in the template dot-products with each STK in the mixture), and for each of the cues, we computed two thresholds: the cue value at the 20th percentile of the cue values within the initial set of 50,000 random STK pairs (the low threshold), and the cue value at the 80th percentile (the high threshold).

There was one experimental condition per cue, and each trial for a condition presented two STK pairs with either high or low values of the cue. The low-value STK pairs were selected to yield cue values that were smaller than the respective low thresholds for each cue. High-value STK pairs were selected to yield a cue value above the high threshold for that cue, while simultaneously having values of all other cues that fell below their respective low thresholds. On each trial a participant heard one low-value and one high-value STK mixture in random order.

During the experiment participants completed 80 trials per condition (320 trials in total). The trials occurred in random order.

#### Experiment 5 - Human agreement with discriminative model decitions

To identify stimulus mixtures classified by the discriminative model as generated by either one or two sources, we first generated 50,000 random pairs of sounds (described for each stimulus type below). We then used the discriminative model to compute the probability of each pair being generated by different sources. We selected the 200 pairs generating the highest probability value and the 200 pairs generating the lowest probability value, and on each trial presented one of each in random order.

The experiment consisted of three blocks, randomly ordered. In each block participants completed 100 trials from each of the following stimulus classes:

1. *STK mixtures*. Each of the two sounds in the mixture corresponded to one of the STKs (drawn from the 10 learned dictionaries).
2. *Modulated noise (spectrotemporal blobs)*.

Each of the two sounds in the mixture was a sample of modulated noise generated using a Gaussian process over the cochleagram. The covariance matrix of the Gaussian process was designed to generate stimuli that were localized and smooth in the cochleagram domain (see below). Each stimulus was randomly drawn as a 40 × 40 pixel shape, subsequently embedded at a random position on the time-frequency plane of the cochleagram (which spanned a time range of [0, 160] ms and a frequency range of [0.02, 8] kHz). Stimuli were thus 160 ms in duration.

The covariance function for each pair of cochleagram pixels *c*_1_, *c*_2_ had the following general form: 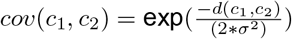, where σ parameter was set to 10. The distance function *d*(*c_1_, c_2_*) had the following form 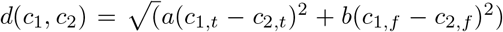, where a and b are parameters controlling the strength of covariance in the temporal and frequency dimension.

To generate a diverse set of stimuli spanning a wide range of spectral and temporal modulation, we used three settings of a and b parameters:

(a) a = 1, b = 1 - these values generated oval-like spectrotemporal shapes

(b) a = 0.1, b = 1 - these values generated temporally elongated, frequency localized, harmonic-like shapes

(c) a = 1, b = 0.1 - these values generated frequency elongated, time-localized, click-like shapes

During stimulus generation, one of these parameter settings was selected randomly with equal probability to generate a sound. We generated a total of 51s sounds which were randomly combined into 50000 pairs.

#### 3.Mixtures of apertured speech

Each of the two sounds in a mixture was generated as follows. We randomly drew 160 ms long excerpts of speech from the TIMIT corpus. Each sample was bandpass filtered and time-windowed to isolate a local patch within the time-frequency plane. We found that this produced stimuli yielding above-chance performance on our task, unlike mixtures of full speech excerpts, which human listeners almost never mistook for a single source. Filtering was performed with a Butterworth filter whose bandwidth was randomly selected to be between 1 and 3 octaves. The lower cutoff of the filter was a random point along the logarithmic frequency axis, constrained to no be more than the Nyquist limit minus the filter bandwidth. After filtering, the waveform of each excerpt was multiplied by a Gaussian window centered at a random position along the excerpt (generated by Matlab function gausswin). The width of the window was controlled by a width parameter proportional to the reciprocal of the standard deviation. The value of the width parameter was randomly drawn from the [1.5, 4] interval with uniform probability.

### Experiment 6 - Streaming of STK sequences

#### Stimulus generation

The stimuli on a trial consisted of a reference sequence paired with a second sequence generated to contain elements that would have either high or low association strength with the elements of the reference sequence. The features within each sequence were spaced further apart in time than those in Experiments 1-5 (75 ms compared to an upper limit of 25 ms in Experiments 1-5).

We generated the STK reference sequence probabilistically using the STK association strength tensor. To generate a sequence, we first chose the first STK in the sequence (each STK was used as the starting STK the same number of times). In the next step we selected a column of the coactivation strength matrix for the first STK corresponding to the desired temporal spacing of the STK to-be-sampled. We used that column to select the next STK in the sequence. To make this choice probabilistic, we transformed this column of coactivation strength values using the softmax transform:

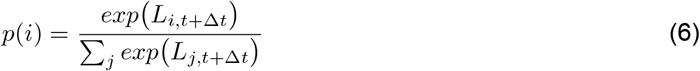

where *L_i,t_* denotes entries of the association strength tensor. The softmax transform generated a discrete probability distribution over the STKs, in which STKs of highest positive association strengths were assigned highest probabilities. STKs with negative association strengths were assigned lowest probabilities. An STK was sampled from the resulting distribution, and this step was iterated to obtain a sequence of the desired length.

The softmax transform is controlled by the “temperature” parameter *β*. If *β* = 0, the probability mass was equal to 0 for all STKs except for the most strongly associated STK, hence the choice was deterministic. For large *β* values (*β* →*∞*), the distribution over STKs became uniform, and all STKs were equally likely regardless of their association strength. The temperature parameter enabled us to generate sequences with varying degrees of randomness.

Sequences of STKs were therefore parametrized by three parameters: the total number of STKs, the temporal spacing of consecutive STKs within a stream, and the temperature parameter controlling the degree of randomness of each stream. All stimuli used here were 4 seconds long (54 STKs). The temporal distance between STKs was set to 75 ms, and the temperature parameter was set to 0.1.

Additionally, to quantify the extent to which the STKs of a sequence should group with each other, we computed the average association strength between consecutive STKs. We refer to this quantity as the “stream coherence”.

For each reference sequence, we generated associated sequences which were either likely or unlikely to co-occur with the reference sequence. We did this by selecting subsets of STKs of either high or low average association strength with the reference sequence. We first computed a weighted average of all columns of the STK tensor used to generate a given sequence. Each column was weighted by the number of occurrences of the corresponding STK in the reference sequence. The resulting average vector had the largest positive values assigned to STKs which were strongly co-activated (on average) with STKs in the stream. The smallest, negative values corresponded to STKs with smallest association strength. We used that average vector to select the 20 most strongly coactivated STKs, or the 20 least strongly coactivated STKs. We then generated sequences in the same way as the reference sequence, only using the selected STKs.

We generated stimuli using a single, randomly chosen STK dictionary. For each of the STKs in the dictionary, we generated 20 random reference sequences with that STK as the first sequence feature, using the procedure described above. For each reference sequence we then generated a co-occurring sequence and a non-co-occurring sequence and added them to the reference sequence, creating two mixtures. This resulted in 20×80=1600 co-occurring sequence mixtures and 1600 non-co-occurring sequence mixtures. We found empirically that non-co-occurring sequences had smaller average stream coherence than co-occurring sequences. To eliminate this difference we selected only the STK sequence mixtures for which the associated co-occurring or non-co-occurring sequences had a stream coherence falling within the interval [0.9, 1.1]. The final stimulus set consisted of 121 co-occurring sequence mixtures and 246 non-co-occurring sequence mixtures, whose associated sequences had approximately the same coherence on average (1.002 and 0.998, respectively). The average stream coherence of the two types of mixtures differed, as intended (1.55 and 0.05 respectively).

#### Experimental procedure

The experiment consisted of 2 blocks of 50 trials. On each trial a participant heard a 4 second-long mixture of a reference stream with either a co-occurring stream or a non-co-occurring stream. Participants judged whether they heard a single source changing in time, or a mixture of two sources. Participants could listen to the stimuli repeatedly if they desired.

#### Participants

Experiments 1, 3, 4, 5 used the same set of 15 participants (8 female, mean age = 25.5, SD = 11.4) who performed the experiments in random order. To ensure task comprehension and motivation, these participants were selected from a larger group of 26 as those who exceeded 90% correct on the speech and instrument condition of Experiment 1. So that we could also measure their performance on this condition without bias from double-dipping, we selected participants using their performance on the odd-numbered trials from this condition, and then analyzed and displayed the performance of the selected participants for the even-numbered trials.

Experiment 2 used a separate set of 15 participants (4 female,mean age = 35, SD = 12.23). To ensure task comprehension and motivation, these participants were selected from a larger group of 23 as those who exceeded an average performance level of 55% correct across both conditions in the experiment (the inclusion criterion was neutral with respect to the hypothesis that performance would be different for natural and artificial co-occurrence statistics).

Experiment 6 used a separate set of 11 participants (6 female, mean age = 36.8, SD = 18.5).

#### Sample sizes

A power analysis performed on pilot data indicated that 14 participants would be needed to reliably detect above-chance performance at the anticipated levels (80% correct; 1 - β = 0.8, α = 0.05). As described above, we ran a larger number of participants and selected those that performed best on speech mixture discrimination, yielding an N of 15 (11 for Experiment 6, which was somewhat easier than the other experiments).

#### Statistics

t tests were used to test for differences in performance between conditions or for differences from chance levels. There was generally one or two such comparisons per experiment, so no correction for multiple comparisons was employed. A repeated-measures ANOVA was used to test for differences among multiple conditions in Experiment 3. Mauchly’s test indicated that the sphericity assumption was violated, and so we report the Greenhouse-Geisser correction. Data distributions were assumed to be normal and were evaluated as such by eye.

## Acknowledgements

We thank Sophia Dolan, River Grace and Malinda McPherson for help with psychophysical experiments, and the McDermott lab for comments on the manuscript. This material is based upon work supported by the Center for Brains, Minds and Machines (CBMM), funded by NSF STC award CCF-1231216. Work also supported by a McDonnell Scholar Award to JHM and NIH Grant No. 1R01DC014739-01A1.

